# Tumor Necrosis Factor and *Schistosoma mansoni* egg antigen Omega-1 shape distinct aspects of the early egg-induced granulomatous response

**DOI:** 10.1101/2020.09.29.318105

**Authors:** Kevin K. Takaki, Francisco J. Roca, Gabriele Schramm, Ruud H. P. Wilbers, Wannaporn Ittiprasert, Paul J. Brindley, Gabriel Rinaldi, Matthew Berriman, Lalita Ramakrishnan, Antonio J. Pagán

## Abstract

Infections by schistosomes result in granulomatous lesions around parasite eggs entrapped within the host tissues. The host and parasite determinants of the *Schistosoma mansoni* egg-induced granulomatous response are areas of active investigation. Some studies in mice implicate Tumor Necrosis Factor (TNF) produced in response to the infection whereas others fail to find a role for it. In addition, in the mouse model, the *S. mansoni* secreted egg antigen omega-1 is found to induce granulomas but the underlying mechanism remains unknown. We have recently developed the zebrafish larva as a model to study macrophage recruitment and granuloma formation in response to *Schistosoma mansoni* eggs. Here we use this model to investigate the mechanisms by which TNF and omega-1 shape the early granulomatous response. We find that TNF, specifically signaling through TNF receptor 1, is not required for macrophage recruitment to the egg and granuloma initiation but does mediate granuloma enlargement. In contrast, omega-1 mediates initial macrophage recruitment, with this chemotactic activity being dependent on its RNase activity. Our findings further the understanding of the role of these host- and parasite-derived factors and show that they impact distinct facets of the granulomatous response to the schistosome egg.

## Introduction

Schistosomiasis is a major granulomatous disease, caused by parasitic flatworms of the genus *Schistosoma* with *Schistosoma mansoni* being the most widespread agent of the disease (McManus et al., 2018). The events of *Schistosoma* egg-induced granulomas have been deduced mainly from histological assessments of human clinical samples and the use of experimental mammalian models (Cheever et al., 2002; Hutchison, 1928). We have recently reported the use of the optically transparent and genetically tractable zebrafish larva as a model to study early macrophage recruitment and granuloma formation in response to *S. mansoni* eggs (Takaki et al., 2020)(Cell Host & Microbe, accepted). Because the zebrafish larva lacks adaptive immunity during their first few weeks of development, this model can be used to dissect mechanisms in the sole context of innate immunity (Davis et al., 2002; Takaki et al., 2013; Takaki et al., 2020). We found that while epithelioid granulomas form rapidly around mature eggs, immature eggs fail to provoke granulomas, consistent with the mature stage-specific secretion of antigens and their function to induce granuloma formation in mammalian models (Ashton et al., 2001; Boros and Warren, 1970; Chiu and Chensue, 2002; Jurberg et al., 2009; Lichtenberg, 1964; Schramm et al., 2006; Takaki et al., 2020).

In the zebrafish, we can additionally examine macrophage recruitment within hours of implantation, and find that whereas injections of schistosome soluble egg antigen (SEA) obtained from mature eggs induce early macrophage recruitment, implantation of immature eggs fail to do so (Takaki et al., 2020). Together these findings both validate the zebrafish model to study *S. mansoni* egg-induced granuloma formation and reveal new insights into the underlying molecular mechanisms (Takaki et al., 2020).

In mice, the cytokine Tumor Necrosis Factor (TNF) and the *S. mansoni* secreted antigen omega-1 have been identified as host and parasite factors, respectively, that promote granuloma formation around the egg (Amiri et al., 1992; Chensue et al., 1994; Chensue et al., 1995; Hagen et al., 2014; Ittiprasert et al., 2019). However, the role of TNF remains controversial and the mechanism by which omega-1 exerts its role is unresolved. In this work, we use the zebrafish model to explore their roles in macrophage recruitment and innate granuloma formation.

## Materials and Methods

### Ethics Statement

All animal experiments were conducted in compliance with guidelines from the UK Home Office and approved by the Wellcome Sanger Institute (WSI) Animal Welfare and Ethical Review Body (AWERB).

### Zebrafish Husbandry

All zebrafish lines were maintained on a recirculating aquaculture system with a 14 hour light - 10 hour dark cycle. Fish were fed dry food and brine shrimp twice a day. Zebrafish embryos were housed in fish water (reverse osmosis water containing 0.18 g/l Instant Ocean) at 28.5°C. Embryos were maintained in 0.25 μg/ml methylene blue from collection to 1 day post-fertilization (dpf). At 24 hours post-fertilization 0.003% PTU (1-phenyl-2-thiourea, Sigma) was added to prevent pigmentation.

### Generation of the TNFR1 mutant and its usage

The zebrafish TNFR1 mutant (*tnfrsf1a*^*rr19*^) was generated using CRISPR Cas9 technology, targeting the sequence TGGTGGAAACAAGACTATGAA of the third exon of the gene (ENSG00000067182) using a T7 promoter-generated guide RNA. Sequencing verified the mutation as a 25 bp deletion (ATGAAGGGAAATTGTCTTGAAAATG) and 6 bp insertion (TGGTGG), resulting in a frame shift and introduction of a premature stop codon soon after the start codon. HRM genotyping was performed using the TNFR1-HRM1-forward and reverse primer set (5’-GTTCCCCACAGGTTCTAACCAG-3’ and 5’-CTTGATGGCATTTATCACAGCAGA-3’, respectively). TNFR1 heterozygotes in the macrophage reporter background, *Tg(mpeg1:YFP)*^*w200*^ (Roca and Ramakrishnan, 2013), were incrossed, genotyped, and sorted as fluorescence-positive, homozygous TNFR1 mutants or WT siblings. Homozygous TNFR1 mutants or WT siblings were then incrossed to generate larvae for experiments.

### Soluble Egg Antigens, WT and RNase mutant recombinant Omega-1

For preparation of SEA, eggs were isolated from *S. mansoni*-infected hamsters as previously described (Schramm et al., 2018), and then homogenized in PBS, pH 7.5, using a sterile glass homogenizer. The homogenate was then centrifuged at 21 krcf for 20 minutes. Supernatants were pooled and then dialyzed overnight in PBS using a 3.5 kDa molecular weight cutoff dialyzer. Sample was then centrifuged at 21 krcf for 20 minutes, and supernatant (SEA) was aliquoted and stored at −80°C. SEA was quantified for protein concentration using the Micro-BCA assay (Pierce, 23225), and quality controlled by SDS-PAGE and western blotting against the *S. mansoni* antigens, omega-1, alpha-1, and kappa-5. Quality control for low LPS content was performed using the Chromo-LAL assay (Associate of Cape Cod, Inc., C0031-5). SEA from WT and corresponding omega-1 knockout eggs were injected at 1 ng per hindbrain ventricle. For comparison of SEA and plant-expressed omega-1, SEA was injected at 2 ng per hindbrain ventricle (1.5 nL injection of 1.4 mg/mL SEA), and plant-expressed omega-1 with LeX glycans (Wilbers et al., 2017) was injected at 0.02 ng per hindbrain ventricle, the relative concentration of omega-1 present in SEA (G. Schramm, personal communication). For DEPC inactivation of plant-expressed omega-1, 1 μL of 0.07 M DEPC (1/100 dilution of Sigma, D5758) was added to 5 μL of 1.5 mg/mL omega-1 (12 mM final concentration of DEPC), and then incubated for 1 hour at 37°C. Because the small volume of protein did not allow for ultrafiltration and requantification of protein, the sample was simply diluted 1/100 in PBS and then 0.02 ng of protein injected into the hindbrain ventricle. For comparison, control sample was incubated at 37°C (without DEPC-treatment) and then diluted 1/100 in PBS. Because the HEK-expressed WT and RNase mutant omega-1 (H58F) lack the native-like LeX glycans in plant-expressed and natural omega-1 (Everts et al., 2012; Everts et al., 2009), they were injected at a 5-fold higher concentration of 0.1 ng per hindbrain ventricle. All hindbrain injections of antigens were assayed at 6 hours post-injection.

### Hindbrain injection of antigens

Hindbrain injections were performed as previously described (Takaki et al., 2013) using 2 ng of WT or Δω1 SEA, 0.02 ng of plant-expressed omega-1 untreated or DEPC-treated, or 0.1 ng of HEK-expressed WT or RNase mutant omega-1.

### Hindbrain implantation of eggs

Schistosome eggs were individually implanted into the zebrafish hindbrain ventricle as previously described (Takaki et al., 2020)(Takaki., Cell Host and Microbe, accepted). Briefly, an incision was made into the zebrafish using a microinjection needle, after which an individual egg was passed though the incision and implanted into the hindbrain ventricle.

### Bacterial infections and quantification of infection burden

Bacterial infections and quantification of infection burden was performed as previously described (Takaki et al., 2013). Briefly, 75 CFU *Mycobacterium marinum* was microinjected into the caudal vein of zebrafish larvae at 36 hours post-fertilization. At 4 days post-infection larvae were imaged by inverted fluorescence microscopy and bacterial fluorescence quantified from images.

### Confocal Microscopy

Zebrafish were anesthetized in fish water containing tricaine and then and mounted onto optical bottom plates (MatTek Corporation, P06G-1.5-20-F) in 1% low melting point agarose (Invitrogen, 16520-100) as previously described (Takaki et al., 2013). Microscopy was performed using a Nikon A1 confocal laser scanning confocal microscopy with a 20x Plan Apo 0.75 NA objective and a Galvano scanner, acquiring 30-80 μm z-stacks with 2-3 μm z-step intervals. Timelapse microscopy was performed at physiological temperature using a heat chamber set to 28°C (Okolab) with an acquisition interval of 3 minutes.

### Granuloma quantification

Confocal images were used for quantifying the number of macrophages in contact with the egg, and subsequent classification of the immune response. Granuloma size was quantified by fluorescence analysis of confocal z-stacks which were flattened, and then fluorescent macrophages comprising the granuloma area was measured by fluorescent pixel counts (FPC) (Takaki et al., 2013).

### Quantification of macrophage recruitment

Quantification of macrophage recruitment was performed by counting the number of fluorescently labeled macrophages within the hindbrain ventricle by fluorescence microscopy. Experiments quantifying macrophage recruitment following injection of egg antigens, utilized *Tg(mpeg1:Brainbow)*^*w201*^ larvae (Pagan et al., 2015).

## Results

### TNF signaling through TNF Receptor 1 promotes macrophage recruitment to nascent *S. mansoni* egg-induced granulomas but is dispensable for initial macrophage recruitment to the eggs

The role of TNF in *S. mansoni* egg-induced granulomas remains unresolved after two decades of studies in the murine model of schistosomiasis. Early findings showed that *S. mansoni*-infected SCID mice were deficient in both granuloma formation and egg extrusion, phenotypes which was rescued by recombinant TNF and activated T cell medium, but not by TNF-depleted T cell medium (Amiri et al., 1992). These findings suggested a role for TNF in granuloma formation and egg excretion (Amiri et al., 1992). However, subsequent work from this group found that TNF knockout mice did not have a defect in granuloma formation (Davies et al., 2004). Mice lacking both receptors through which TNF signals did exhibit a mild granuloma deficit, leading the authors to propose that it might be due to a defect in signaling of the ligand lymphotoxin (Davies et al., 2004). However, this would not explain their earlier findings that exogenous TNF rescued granuloma formation in SCID mice (Amiri et al., 1992). Meanwhile, a different group reported that TNF did not rescue granuloma formation in SCID mice (Cheever et al., 1999). Additionally, *S. mansoni*-infected SCID mice displayed normal levels of TNF expression, suggesting that other cells may be the major source of TNF during the infection (Cheever et al., 1999). It has been suggested that Ly6C^hi^ monocytes, which are known to express TNF in response to the schistosome egg, might be the innate source of TNF (Nascimento et al., 2014).

To delineate the role of TNF in macrophage recruitment and granuloma formation around S. mansoni eggs, we used a TNFR1 zebrafish mutant created by CRISPR-Cas technology (see Methods). We first confirmed that the lack of TNFR1 signaling rendered zebrafish larvae susceptible to *Mycobacterium marinum* infection, consistent to our previous findings using TNFR1 morpholino (Clay et al., 2008)(**Figure S1**). Next, we implanted the Hindbrain Ventricle (HBV) of wildtype and TNFR1 mutant larvae with *S. mansoni* eggs and evaluated granuloma formation after five days (Figure 1A-C). We have recently categorized early macrophage recruitment and granuloma formation based on the number and characteristics of macrophages in contact with the egg: Minimal recruitment, 0-6 macrophages; Macrophages recruited, >6 macrophages; Granulomas, confluent epithelioid macrophage aggregates (Takaki et al., 2020)(Cell Host & Microbe, accepted). At 5 days post-implantation of the eggs, the TNFR1 mutants had similar macrophage responses to wildtype animals with ∼50% of the animals forming epithelioid granulomas in each group (**Figure 1B**). However, we found that the TNFR1 deficient granulomas were significantly smaller than wildtype granulomas, with the mean granuloma size being 62% smaller than in wildtype (**Figure 1C** and **1D**). We also noted that the TNFR1 mutant granulomas, though smaller, showed a characteristic epithelioid morphology with confluent macrophages and loss of intercellular boundaries. This finding suggest that epithelioid transformation may also be independent of TNFR1 signaling (Takaki et al., 2020)(Cell Host & Microbe, accepted)(**Figure 1D**). Because the *S. mansoni* granuloma is comprised solely of macrophages at this early stage (Takaki et al., 2020)(Cell Host & Microbe, accepted), our findings imply that TNFR1 signaling would promote macrophage recruitment to a nascent granuloma around the egg. In the zebrafish model, we can also examine the initiation of macrophage recruitment to *S. mansoni* eggs within hours of implantation (Takaki et al., 2020)(Cell Host & Microbe, accepted). However, we found that TNFR1 signaling is not required for initiation of macrophage recruitment (**Figure 1E**).

**Figure 1.**
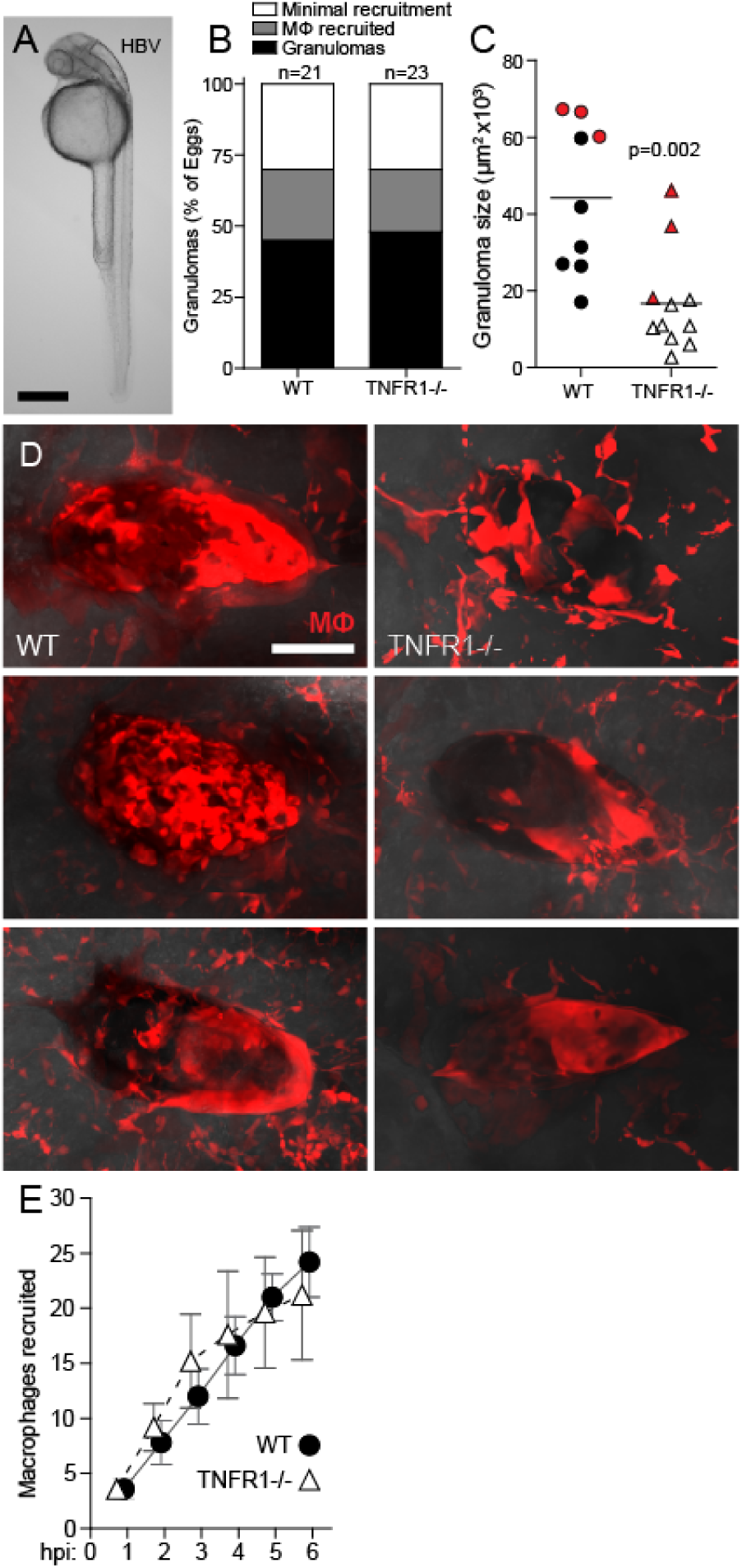
TNF affects late-stage granuloma formation. Comparison of macrophage recruitment and granuloma formation in WT and TNFR1 mutant zebrafish larvae following implantation with a single schistosome egg into their hindbrain ventricle. (**A**) Zebrafish larva at 36 hours post-fertilization with the hindbrain ventricle (HBV) site of injection and implantation outlined. Scale bar, 300 μm. (**B**-**D**) Granuloma formation at 5 days post-implantation. (**B**) Percent of animals with; granuloma formation (confluent epithelioid macrophage aggregates), macrophages recruited (>6 macrophages in contact with the egg), or minimal recruitment (0-6 macrophages in contact with egg) (Takaki et al., 2020)(Cell Host & Microbe, accepted). (**C**) Granuloma size and (**D**) images, with each image from top to bottom corresponding with each red data point, top to bottom, respectively. Scale bar, 50 μm. Horizontal bars in (C), means. Statistics, Student’s *t-*test. (**E**) Mean macrophage recruitment kinetics during the first 6 hours post-implantation. Error bars, SEM. Sample size, n=5 animals per group.

Together, these results show that TNF signaling through TNFR1 is required specifically for macrophage recruitment after the initial macrophages reach the egg through other signal(s). Thereby, TNF mediates granuloma enlargement rather than granuloma initiation. Furthermore, TNF is not required for epithelioid transformation. Finally, TNF plays a role in the granulomatous response in the sole context of innate immunity.

### *S. mansoni* omega-1 promotes initial macrophage recruitment to the egg through its RNase activity

Next, we wanted to probe the parasite determinants that induce granuloma formation. In recent work we found that immature *S. mansoni* eggs invoked neither granuloma formation nor even initial macrophage recruitment, indicating that mature egg antigens were required for the first macrophages to be recruited to the egg (Takaki et al., 2020)(Cell Host & Microbe, accepted). Mature eggs express a variety of antigens (Ashton et al., 2001; Cass et al., 2007; Dunne et al., 1981), of which omega-1 is known to be the major contributor to granuloma formation, as knockdown of its expression leads to greatly diminished granuloma formation around eggs (Hagen et al., 2014; Ittiprasert et al., 2019). Omega-1 is an RNase involved in several processes. In dendritic cells, it inhibits protein synthesis, alters cell morphology, induces IL-33 expression, and reduces conjugation affinity with T cells (Everts et al., 2012; Everts et al., 2009; Fitzsimmons et al., 2005; Steinfelder et al., 2009). If and how this leads to granuloma formation is not known. However, it is well-established that its RNase activity is essential for inducing the Th2 polarization of granulomas (Everts et al., 2012; Everts et al., 2009; Fitzsimmons et al., 2005; Steinfelder et al., 2009). This in turn induces expression of IL-4 and IL-13, known egg-induced host factors that can mediate granuloma formation (Cheever et al., 1999; Fallon et al., 2000; Jankovic et al., 1999). Additionally, omega-1 is a major hepatotoxin (Abdulla et al., 2011; Dunne et al., 1991; Dunne et al., 1981), and it has been proposed that the granuloma itself would prevent the cytotoxic effects of this egg antigen on the host liver.

Our attempts to test the role of omega-1 by implanting omega-1 knockout (KO) eggs (Ittiprasert et al., 2019) into the larvae failed, as the genetically modified eggs did not survive shipment. As an alternative approach, we tested if the SEA obtained from omega-1 KO eggs could recruited macrophages. We examined macrophage recruitment 6 hours post-injection of the SEA into the hindbrain ventricle (**Figure 1A**). Omega-1-deficient SEA recruited macrophages similar to wildtype SEA (**Figure 2A**). The omega-1 deficient SEA retains ∼20% of the omega-1 RNase activity (not shown), suggesting that even though reduced compared to wild type eggs, it is may still be sufficient for macrophage recruitment (Ittiprasert et al., 2019). Alternatively, the omega-1 activity may be redundant with other SEA components (Kaisar et al., 2018). To investigate these hypotheses, we used a recombinant omega-1 that contain the native-like LeX glycosylation, which is important for its uptake by dendritic cells and subsequent Th2-polarization (Everts et al., 2012; Wilbers et al., 2017). Injection of 0.02 ng of omega-1, the approximate amount of omega-1 in the corresponding SEA injections (Cell Host & Microbe, accepted, G. Schramm, personal communication), induced macrophage recruitment, although less than SEA, consistent with other components inducing macrophage recruitment (**Figure 2B and 2C**).

**Figure 2.**
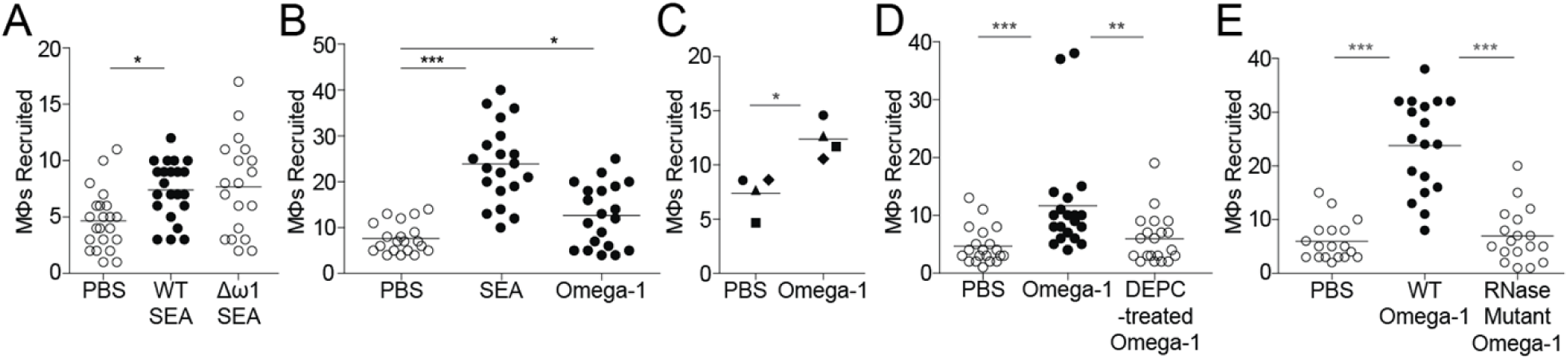
Omega-1 recruits macrophages via its RNase activity. Macrophage recruitment at 6 hours post-injection (hpi) with egg antigens. (**A**) Macrophages recruited to SEA from WT or omega-1 knockout eggs (Δω1). (**B**) Macrophages recruited to SEA or omega-1. (**C**) Mean macrophage recruitment to omega-1 for each of four experiments. Individual experiments represented with unique symbols; triangles and squares represent means of panels B and D. (**D**) Macrophages recruited to omega-1 or DEPC-treated omega-1. (**E**) Macrophages recruited to WT or omega-1 mutant. All omega-1 injections were performed using 0.02 ng of plant-expressed omega-1, with the exception of (**E**) which used HEK-expressed WT or mutant omega-1 injected at a 5-fold higher concentration of 0.1 ng to compensate for lack of LeX glycosylation in plant-expressed and natural omega-1. Statistics, ANOVA with Dunnett’s post-test comparing all samples to PBS (**B**) or WT omega-1 (**A**,**E**); (**D**) non-parametric ANOVA with Dunn’s post-test comparing all samples to omega-1; (**C**) paired *t*-test. All horizontal bars, means.

Next, we asked if omega-1-associated recruitment of macrophages is dependent on its RNase activity. The inhibition of RNase activity in the recombinant omega-1 with diethyl pyrocarbonate (DEPC) (Steinfelder et al., 2009), led to loss of macrophage-recruiting activity (**Figure 2D**). Because DEPC inhibits RNase function through covalent binding to the essential histidine in the catalytic domains of RNase, one caveat is that it can create off-target modifications to the protein structure and function through binding to other histidine residues as well as, to a lesser extent, tyrosine, lysine, and cysteine (Wolf et al., 1970). To validate our findings, we used recombinant omega-1 mutant lacking RNase activity, with a phenylalanine substitution of the essential histidine of the catalytic domain (Everts et al., 2012; Irie and Ohgi, 2001). As expected, the omega-1 mutant failed to recruit macrophages (**Figure 2E**). These findings confirmed that the omega-1 macrophage chemotactic activity is mediated through its RNase activity (**Figure 2B**).

## Discussion

This study reinforces the use of the zebrafish model to study molecular pathways involved in *S. mansoni*-egg-induced granuloma formation. Particularly, it provides new insights on host and parasite factors modulating this critical process that drives the pathology associated with schistosomiasis.

We demonstrate that the cytokine TNF is required for granuloma enlargement but not initiation, in agreement with previous observations in the mouse (Amiri et al., 1992; Chensue et al., 1994; Chensue et al., 1995; Ehlers and Schaible, 2012). Further, we show that TNF is dispensable for the first wave of macrophage recruitment to the egg. These findings are consistent with TNF not being a direct chemotactic agent, but mediating cell recruitment through interactions with other cells that, in turn, synthesize macrophage chemokines (Kalliolias and Ivashkiv, 2016; Mukaida et al., 2011). SEA is known to induce the expression of TNF (Chensue et al., 1994; Chiu et al., 2004; Nascimento et al., 2014), therefore, we reason that it might be only after granuloma initiation, at which point significant numbers of macrophages are in contact with the egg, that TNF is produced above the threshold to induce these chemokines. In addition, the close cell-to-cell contacts following the initiation of granuloma formation and epithelioid transformation may be vital; if TNF is acting in both an autocrine and paracrine manner, then the cell-to-cell interaction would allow for maximal signal exchange between cells, the optimal amplification of this signal and subsequent expression of chemokines (Blasi et al., 1994; Caldwell et al., 2014). Epithelioid transformation is primarily associated with Th2-polarized immune responses involving IL-4/IL-13, expression of which can occur in the context of innate immunity alone (Bottiglione et al., 2020; Chiu et al., 2004; Mitre et al., 2004; Pagan and Ramakrishnan, 2018). Therefore, it is not surprising to observe epithelioid transformation in the absence of TNF. Chronic mTORC1 signaling, which does not require adaptive immunity, can also induce epithelioid transformation (Linke et al., 2017).

We have recently shown that *S. mansoni* eggs, upon reaching maturity, induce granuloma formation that benefits the parasite by extruding the egg into the environment (Takaki et al., 2020) (Cell Host & Microbe, accepted). This would be achieved by mature egg stage-dependent secretion of antigens such as omega-1 (Ashton et al., 2001; Schramm et al., 2006). Here, we show that recombinant omega-1 recruited macrophages rapidly, similar to SEA. Our finding supports the hypothesis that omega-1 is sufficient yet dispensable for early macrophage recruitment. This may have parallels in observations regarding its role in granuloma formation; omega-1 knockdown eggs form granulomas in the mouse, albeit smaller ones, suggesting other egg antigens such as IPSE could contribute to this process (Hagen et al., 2014; Ittiprasert et al., 2019).

In addition, we have demonstrated that the omega-1 RNase activity is required for macrophage recruitment. This finding indicates that omega-1 does not act directly as a chemoattractant, and that recruitment must be mediated through downstream effects stemming from its RNase activity. Prior work has shown that its RNase activity mediates Th2 polarization through inhibition of protein synthesis in dendritic cells (Everts et al., 2012; Everts et al., 2009; Steinfelder et al., 2009). In the context of the 6-hour recruitment assay performed herein, we speculate that the protein is taken up by epithelial cells that line the hindbrain ventricle cavity, perturbing cellular homeostasis by an RNase-induced inhibition of protein synthesis and in turn, inducing cell stress signals which would trigger macrophage recruitment

As for tuberculous granulomas (Pagan and Ramakrishnan, 2018; Ramakrishnan, 2012), we expect this report will stimulate the use of this facile model to dissect mechanisms underlying the genesis of schistosome egg-induced granulomas, main drivers of schistosomiasis pathogenesis and transmission.

## Acknowledgements

This work was supported by Wellcome Trust core-funding support to the Wellcome Sanger Institute (award number 206194) (GR, MB) and NIH MERIT award (R37 AI054503) and a Wellcome Trust Principal Research Fellowship (LR). These studies were additionally supported the Wellcome Trust Strategic Award number 107475/Z/15/Z, and the NIAID Schistosomiasis Resource Center for distribution through BEI Resources, NIH-NIAID Contract HHSN272201000005I (PJB, WI).

## Contributions

Conceived and designed experiments: KKT LR AJP. Performed experiments: KKT. Analyzed data: LR KKT AJP. Contributed reagents/materials/analysis tools: FJR GS RHPW WI PJB GR MB. Wrote paper: KKT, LR. Edited paper: FJR AJP RHPW WI PJB GR MB.

---

**Figure S1.**
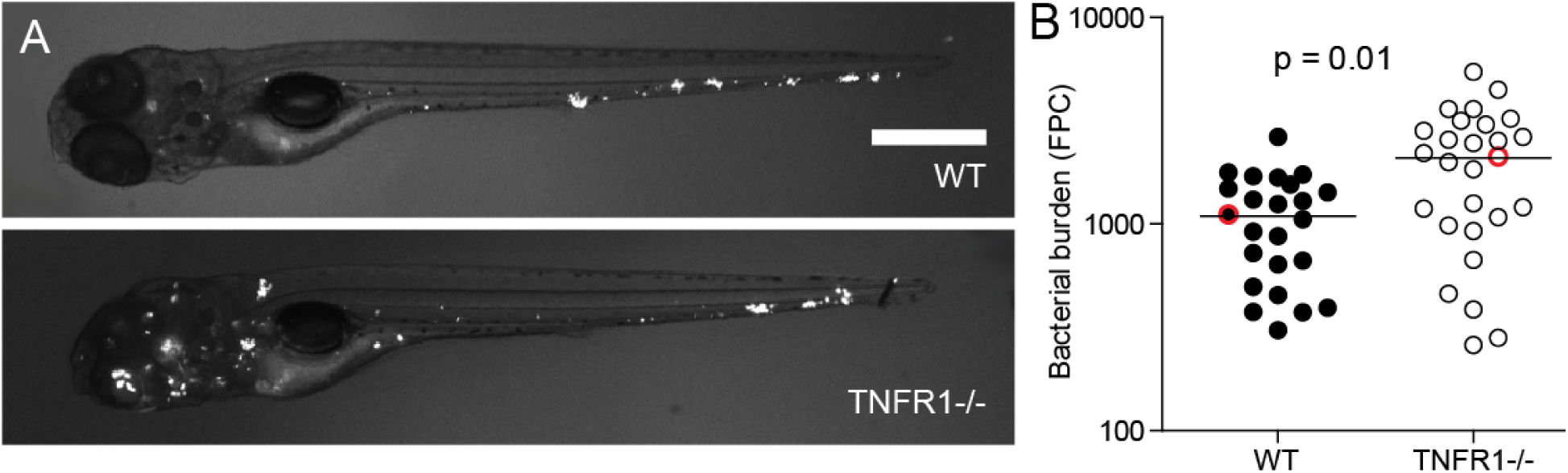
TNFR1 mutant zebrafish larvae have increased infection burden. WT and TNFR1 mutant zebrafish larvae were systemically infected at 36 hours post-fertilization via caudal vein injection with 75 CFU *Mycobacterium marinum*, and then imaged at 4 days post-infection for bacterial burden. (**A**) The two animals closest to the mean. Scale bar, 300 μm. (**B**) Quantification of bacterial burden, with the two red data points corresponding to the animals in (A). Horizontal bar, means. Statistics, Student’s t test. FPC: fluorescent pixel counts.

## References

Abdulla, M.H., Lim, K.C., McKerrow, J.H., and Caffrey, C.R. (2011). Proteomic identification of IPSE/alpha-1 as a major hepatotoxin secreted by Schistosoma mansoni eggs. PLoS Negl Trop Dis 5, e1368.

Amiri, P., Locksley, R.M., Parslow, T.G., Sadick, M., Rector, E., Ritter, D., and McKerrow, J.H. (1992). Tumour necrosis factor alpha restores granulomas and induces parasite egg-laying in schistosome-infected SCID mice. Nature 356, 604–607.

Ashton, P.D., Harrop, R., Shah, B., and Wilson, R.A. (2001). The schistosome egg: development and secretions. Parasitology 122, 329–338.

Blasi, E., Pitzurra, L., Bartoli, A., Puliti, M., and Bistoni, F. (1994). Tumor necrosis factor as an autocrine and paracrine signal controlling the macrophage secretory response to Candida albicans. Infect Immun 62, 1199–1206.

Boros, D.L., and Warren, K.S. (1970). Delayed hypersensitivity-type granuloma formation and dermal reaction induced and elicited by a soluble factor isolated from Schistosoma mansoni eggs. J Exp Med 132, 488–507.

Bottiglione, F., Dee, C.T., Lea, R., Zeef, L.A.H., Badrock, A.P., Wane, M., Bugeon, L., Dallman, M.J., Allen, J.E., and Hurlstone, A.F.L. (2020). Zebrafish IL-4-like Cytokines and IL-10 Suppress Inflammation but Only IL-10 Is Essential for Gill Homeostasis. J Immunol 205, 994–1008.

Caldwell, A.B., Cheng, Z., Vargas, J.D., Birnbaum, H.A., and Hoffmann, A. (2014). Network dynamics determine the autocrine and paracrine signaling functions of TNF. Genes Dev 28, 2120–2133.

Cass, C.L., Johnson, J.R., Califf, L.L., Xu, T., Hernandez, H.J., Stadecker, M.J., Yates, J.R., 3rd, and Williams, D.L. (2007). Proteomic analysis of Schistosoma mansoni egg secretions. Mol Biochem Parasitol 155, 84–93.

Cheever, A.W., Lenzi, J.A., Lenzi, H.L., and Andrade, Z.A. (2002). Experimental models of Schistosoma mansoni infection. Mem Inst Oswaldo Cruz 97, 917–940.

Cheever, A.W., Poindexter, R.W., and Wynn, T.A. (1999). Egg laying is delayed but worm fecundity is normal in SCID mice infected with Schistosoma japonicum and S. mansoni with or without recombinant tumor necrosis factor alpha treatment. Infect Immun 67, 2201–2208.

Chensue, S.W., Warmington, K., Ruth, J., Lincoln, P., Kuo, M.C., and Kunkel, S.L. (1994). Cytokine responses during mycobacterial and schistosomal antigen-induced pulmonary granuloma formation. Production of Th1 and Th2 cytokines and relative contribution of tumor necrosis factor. Am J Pathol 145, 1105–1113.

Chensue, S.W., Warmington, K.S., Ruth, J.H., Lincoln, P., and Kunkel, S.L. (1995). Cytokine function during mycobacterial and schistosomal antigen-induced pulmonary granuloma formation. Local and regional participation of IFN-gamma, IL-10, and TNF. J Immunol 154, 5969–5976.

Chiu, B.C., and Chensue, S.W. (2002). Chemokine responses in schistosomal antigen-elicited granuloma formation. Parasite Immunol 24, 285–294.

Chiu, B.C., Freeman, C.M., Stolberg, V.R., Hu, J.S., Komuniecki, E., and Chensue, S.W. (2004). The innate pulmonary granuloma: characterization and demonstration of dendritic cell recruitment and function. Am J Pathol 164, 1021–1030.

Clay, H., Volkman, H.E., and Ramakrishnan, L. (2008). Tumor necrosis factor signaling mediates resistance to mycobacteria by inhibiting bacterial growth and macrophage death. Immunity 29, 283–294.

Davies, S.J., Lim, K.C., Blank, R.B., Kim, J.H., Lucas, K.D., Hernandez, D.C., Sedgwick, J.D., and McKerrow, J.H. (2004). Involvement of TNF in limiting liver pathology and promoting parasite survival during schistosome infection. Int J Parasitol 34, 27–36.

Davis, J.M., Clay, H., Lewis, J.L., Ghori, N., Herbomel, P., and Ramakrishnan, L. (2002). Real-time visualization of mycobacterium-macrophage interactions leading to initiation of granuloma formation in zebrafish embryos. Immunity 17, 693–702.

Dunne, D.W., Jones, F.M., and Doenhoff, M.J. (1991). The purification, characterization, serological activity and hepatotoxic properties of two cationic glycoproteins (alpha 1 and omega 1) from Schistosoma mansoni eggs. Parasitology 103 Pt 2, 225–236.

Dunne, D.W., Lucas, S., Bickle, Q., Pearson, S., Madgwick, L., Bain, J., and Doenhoff, M.J. (1981). Identification and partial purification of an antigen (omega 1) from Schistosoma mansoni eggs which is putatively hepatotoxic in T-cell deprived mice. Trans R Soc Trop Med Hyg 75, 54–71.

Ehlers, S., and Schaible, U.E. (2012). The granuloma in tuberculosis: dynamics of a host-pathogen collusion. Front Immunol 3, 411.

Everts, B., Hussaarts, L., Driessen, N.N., Meevissen, M.H., Schramm, G., van der Ham, A.J., van der Hoeven, B., Scholzen, T., Burgdorf, S., Mohrs, M., et al. (2012). Schistosome-derived omega-1 drives Th2 polarization by suppressing protein synthesis following internalization by the mannose receptor. J Exp Med 209, 1753-1767, S1751.

Everts, B., Perona-Wright, G., Smits, H.H., Hokke, C.H., van der Ham, A.J., Fitzsimmons, C.M., Doenhoff, M.J., van der Bosch, J., Mohrs, K., Haas, H., et al. (2009). Omega-1, a glycoprotein secreted by Schistosoma mansoni eggs, drives Th2 responses. J Exp Med 206, 1673–1680.

Fallon, P.G., Richardson, E.J., McKenzie, G.J., and McKenzie, A.N. (2000). Schistosome infection of transgenic mice defines distinct and contrasting pathogenic roles for IL-4 and IL-13: IL-13 is a profibrotic agent. J Immunol 164, 2585–2591.

Fitzsimmons, C.M., Schramm, G., Jones, F.M., Chalmers, I.W., Hoffmann, K.F., Grevelding, C.G., Wuhrer, M., Hokke, C.H., Haas, H., Doenhoff, M.J., et al. (2005). Molecular characterization of omega-1: a hepatotoxic ribonuclease from Schistosoma mansoni eggs. Mol Biochem Parasitol 144, 123–127.

Hagen, J., Young, N.D., Every, A.L., Pagel, C.N., Schnoeller, C., Scheerlinck, J.P., Gasser, R.B., and Kalinna, B.H. (2014). Omega-1 knockdown in Schistosoma mansoni eggs by lentivirus transduction reduces granuloma size in vivo. Nat Commun 5, 5375.

Hutchison, H.S. (1928). The Pathology of Bilharziasis. Am J Pathol 4, 1-16 11.

Irie, M., and Ohgi, K. (2001). Ribonuclease T2. Methods Enzymol 341, 42–55.

Ittiprasert, W., Mann, V.H., Karinshak, S.E., Coghlan, A., Rinaldi, G., Sankaranarayanan, G., Chaidee, A., Tanno, T., Kumkhaek, C., Prangtaworn, P., et al. (2019). Programmed genome editing of the omega-1 ribonuclease of the blood fluke, Schistosoma mansoni. Elife 8.

Jankovic, D., Kullberg, M.C., Noben-Trauth, N., Caspar, P., Ward, J.M., Cheever, A.W., Paul, W.E., and Sher, A. (1999). Schistosome-infected IL-4 receptor knockout (KO) mice, in contrast to IL-4 KO mice, fail to develop granulomatous pathology while maintaining the same lymphokine expression profile. J Immunol 163, 337–342.

Jurberg, A.D., Goncalves, T., Costa, T.A., de Mattos, A.C., Pascarelli, B.M., de Manso, P.P., Ribeiro-Alves, M., Pelajo-Machado, M., Peralta, J.M., Coelho, P.M., et al. (2009). The embryonic development of Schistosoma mansoni eggs: proposal for a new staging system. Dev Genes Evol 219, 219–234.

Kaisar, M.M.M., Ritter, M., Del Fresno, C., Jonasdottir, H.S., van der Ham, A.J., Pelgrom, L.R., Schramm, G., Layland, L.E., Sancho, D., Prazeres da Costa, C., et al. (2018). Dectin-1/2-induced autocrine PGE2 signaling licenses dendritic cells to prime Th2 responses. PLoS Biol 16, e2005504.

Kalliolias, G.D., and Ivashkiv, L.B. (2016). TNF biology, pathogenic mechanisms and emerging therapeutic strategies. Nat Rev Rheumatol 12, 49–62.

Lichtenberg (1964). Studies on Granuloma Formation. Iii. Antigen Sequestration and Destruction in the Schistosome Pseudotubercle. Am J Pathol 45, 75–94.

Linke, M., Pham, H.T., Katholnig, K., Schnoller, T., Miller, A., Demel, F., Schutz, B., Rosner, M., Kovacic, B., Sukhbaatar, N., et al. (2017). Chronic signaling via the metabolic checkpoint kinase mTORC1 induces macrophage granuloma formation and marks sarcoidosis progression. Nat Immunol 18, 293–302.

McManus, D.P., Dunne, D.W., Sacko, M., Utzinger, J., Vennervald, B.J., and Zhou, X.N. (2018). Schistosomiasis. Nat Rev Dis Primers 4, 13.

Mitre, E., Taylor, R.T., Kubofcik, J., and Nutman, T.B. (2004). Parasite antigen-driven basophils are a major source of IL-4 in human filarial infections. J Immunol 172, 2439–2445.

Mukaida, N., Sasakki, S., and Popivanova, B.K. (2011). Tumor Necrosis Factor (TNF) and Chemokines in Colitis-Associated Cancer. Cancers (Basel) 3, 2811–2826.

Nascimento, M., Huang, S.C., Smith, A., Everts, B., Lam, W., Bassity, E., Gautier, E.L., Randolph, G.J., and Pearce, E.J. (2014). Ly6Chi monocyte recruitment is responsible for Th2 associated host-protective macrophage accumulation in liver inflammation due to schistosomiasis. PLoS Pathog 10, e1004282.

Pagan, A.J., and Ramakrishnan, L. (2018). The Formation and Function of Granulomas. Annu Rev Immunol 36, 639–665.

Pagan, A.J., Yang, C.T., Cameron, J., Swaim, L.E., Ellett, F., Lieschke, G.J., and Ramakrishnan, L. (2015). Myeloid Growth Factors Promote Resistance to Mycobacterial Infection by Curtailing Granuloma Necrosis through Macrophage Replenishment. Cell Host Microbe 18, 15–26.

Ramakrishnan, L. (2012). Revisiting the role of the granuloma in tuberculosis. Nat Rev Immunol 12, 352–366.

Roca, F.J., and Ramakrishnan, L. (2013). TNF dually mediates resistance and susceptibility to mycobacteria via mitochondrial reactive oxygen species. Cell 153, 521–534.

Schramm, G., Gronow, A., Knobloch, J., Wippersteg, V., Grevelding, C.G., Galle, J., Fuller, H., Stanley, R.G., Chiodini, P.L., Haas, H., et al. (2006). IPSE/alpha-1: a major immunogenic component secreted from Schistosoma mansoni eggs. Mol Biochem Parasitol 147, 9–19.

Schramm, G., Suwandi, A., Galeev, A., Sharma, S., Braun, J., Claes, A.K., Braubach, P., and Grassl, G.A. (2018). Schistosome Eggs Impair Protective Th1/Th17 Immune Responses Against Salmonella Infection. Front Immunol 9, 2614.

Steinfelder, S., Andersen, J.F., Cannons, J.L., Feng, C.G., Joshi, M., Dwyer, D., Caspar, P., Schwartzberg, P.L., Sher, A., and Jankovic, D. (2009). The major component in schistosome eggs responsible for conditioning dendritic cells for Th2 polarization is a T2 ribonuclease (omega-1). J Exp Med 206, 1681–1690.

Takaki, K., Davis, J.M., Winglee, K., and Ramakrishnan, L. (2013). Evaluation of the pathogenesis and treatment of Mycobacterium marinum infection in zebrafish. Nat Protoc 8, 1114–1124.

Takaki, K.K., Rinaldi, G., Berriman, M., Pagán, A.J., and Ramakrishnan, L. (2020). Schistosoma mansoni eggs modulate the timing of granuloma formation to promote transmission. bioRxiv.

Wilbers, R.H., Westerhof, L.B., van Noort, K., Obieglo, K., Driessen, N.N., Everts, B., Gringhuis, S.I., Schramm, G., Goverse, A., Smant, G., et al. (2017). Production and glyco-engineering of immunomodulatory helminth glycoproteins in plants. Sci Rep 7, 45910.

Wolf, B., Lesnaw, J.A., and Reichmann, M.E. (1970). A mechanism of the irreversible inactivation of bovine pancreatic ribonuclease by diethylpyrocarbonate. A general reaction of diethylpyrocarbonate. A general reaction of diethylpyrocarbonate with proteins. Eur J Biochem 13, 519–525.

